# The Dynamics and Energy Landscape of DNA Plectoneme Nucleation

**DOI:** 10.1101/280719

**Authors:** Philipp U. Walker, Willem Vanderlinden, Jan Lipfert

**Affiliations:** Department of Physics, Nanosystems Initiative Munich, and Center for Nanoscience, LMU Munich, Amalienstrasse54, 80799 Munich, Germany; Department of Chemistry, KU Leuven, Celestijnenlaan 200F, 3001 Heverlee, Belgium

## Abstract

DNA buckling is the fundamental step for plectoneme nucleation and supercoil dynamics that are critical in the processing of genetic information. Here we systematically quantify DNA buckling dynamics using high-speed magnetic tweezers. Buckling times are ∼10-100 ms and depend exponentially on both applied force and twist. By deconvolving measured time traces with the instrument response, we reconstruct full 2D extension-twist energy landscapes of the buckling transition that reveal an asymmetry between the pre- and post-buckling states and suggest a highly bend transition state conformation.

## INTRODUCTION

DNA stores genetic information as a linear sequence and consequently needs to be very long. To achieve compaction in the narrow confines of the cell, while providing local accessibility for readout and processing, genome architecture is dynamically controlled [1-4]. Regulation is achieved by organizing DNA into domains, wherein DNA rotational motion is constrained. As a result, the number of links between the two single strands of the double helix, called the linking number *Lk*, is invariant. The topological quantity *LK* partitions into the geometric parameters twist *Tw* and writhe *Wr* [5,6],

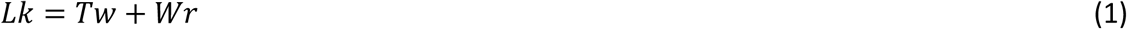

where *Tw* quantifies the torsional deformations and strains of the helix, and *Wr* the coiling of the double helix axis in 3D. Specialized proteins maintain cellular DNA in a supercoiled state, i.e. *LK* deviates from the torsionally relaxed value *LK*^*0*^ *in vivo*. Through topological coupling, the linking difference *ΔLK= LK – LK*^*0*^ affects DNA structure both locally and globally, via changes in twist *ΔTw* and writhe *ΔWr* respectively [3,7,8]. Supercoiled DNA, in general, is both locally untwisted and takes on interwound, plectonemic configurations of the double helix axis. The structure and mechanics of plectonemic DNA has been probed extensively by electron and atomic force microscopy of circular DNA molecules and DNA tethered between a surface and magnetic beads in magnetic tweezers (MT) (**Fig. 1a**) [9-11]. In contrast, the dynamics of plectonemes remain largely unexplored, despite their importance in the context of regulation and long-range communication in the cell [12].

**Figure 1.**
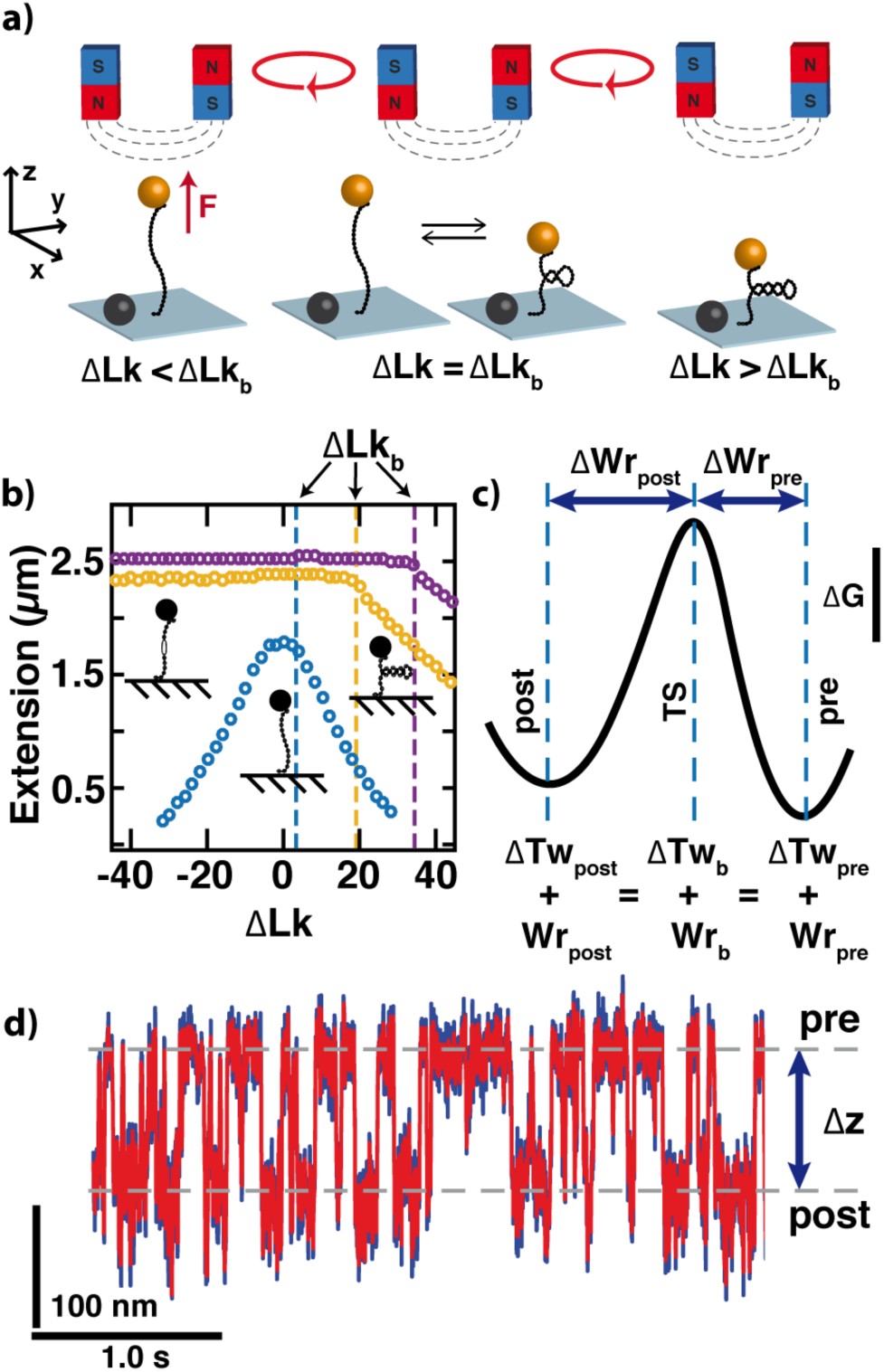
DNA buckling measurements in the MT. (a) Schematic of the MT setup. A DNA molecule is tethered between the surface and a superparamagnetic bead. Magnets exert a force on the bead and rotation of the magnets controls the linking number. Close to the buckling point *ΔLK*_*b*_ the DNA extension jumps between two states. (b) Rotation-extension curves for different forces. At low forces (blue, 0.2 pN) DNA forms plectonemes at positive and negative turns. At higher forces (yellow, 2 pN; purple, 5 pN) plectonemes appear only at positive turns, while DNA melts at negative *ΔLK*. (c) Schematic representation of the buckling energy landscape along the rotational coordinate at *ΔLK*_*b*_. (d) Extension trace close to *ΔLK*_*b*_, depicting the dynamic hopping of the DNA molecule between the pre-buckling and post-nucleation states. *Δz* indicates the jump size (raw data at 1000 Hz, blue; smoothed data at 333 Hz, red).

A first direct measurement of plectonemes dynamics by fluorescent imaging [13] found that, within the time-resolution of the approach (20 ms), a plectoneme can disappear at one site and give rise to the appearance of a new plectoneme separated by several microns along the chain. The dynamics of this process are proposed to be rate-limited by nucleation of the newly formed plectoneme, i.e. by DNA buckling, in particular since removal of the plectoneme has been shown to be very fast [14]. Estimates for the characteristic time scale of DNA buckling range from 30-80 ms in magnetic tweezers (MT) [11,15], to ∼100 ms in an optical torque wrench [16,17], and are, surprisingly, ∼2 order-of-magnitude faster than for double-stranded RNA [15]. However, a direct comparison of the different measurements is complicated as the dynamics depend on ionic strength and applied stretching force. The lack of a precise quantitation of DNA buckling dynamics is at least in part due to the difficulty of observing fast ∼ms time scale transitions using camera based detection.

Here, we have used high-speed magnetic tweezers, to quantify the kinetics of supercoil nucleation under a range of forces, ionic strength, and bead sizes. Using a deconvolution approach, we reconstruct the 2D energy landscape of the buckling transition as a function of DNA extension and twist. We propose a geometric model for the transition state and discuss how local disturbances of the helix can impact the energy landscape of supercoiling.

## RESULTS AND DISCUSSION

### Bead tracking with kHz time-resolution accurately quantifies DNA buckling transitions

In our MT setup, linear ∼7.9 kbp DNA molecules are tethered between the flow cell surface and superparamagnetic beads, via multiple attachment points at both ends to assure torsionally constrained attachment (**Fig. 1a** and **Supplementary Material** [18]). We note that our DNA length approximately corresponds to the size of topological DNA domains *in vivo*, ∼10 kbp [19]. Using external magnets, one can apply precisely calibrated forces [10,20,21] and control the linking number of the DNA. On increasing *ΔLK*at forces < 6 pN [22], DNA undergoes a buckling transition and starts to form plectonemes, causing a decrease in the apparent tether extension with increasing *ΔLK*(**Fig. 1b**). If DNA molecules are held at a fixed *ΔLK*close to the buckling point *ΔLK*_*b*_, the molecules undergo thermally activated transitions between the torsionally strained, but extended pre-nucleation state, and the post-nucleation state, with the first, minimal plectoneme formed (**Fig. 1c**). Here, we draw on improvements in both camera and illumination hardware as well as tracking software [23-25] to study supercoil nucleation by tracking at 1 kHz in MT. Simulations show that our experimental configuration can resolve transitions on the time scale of ∼10 ms or less, with an error of at most 10% (**Fig. 1d** and **Fig. S1** in [18]). In addition, from analysis of experimental extension time traces of torsionally relaxed DNA tethers, we find the characteristic (“corner”) frequencies at 2, 3 and 4 pN to be 98 ± 3 Hz, 161 ± 6 Hz and 230 ± 12 Hz (means and standard errors from 4 independent measurements at each force), respectively, corresponding to characteristic time scales of 10, 6, and 4 ms, again suggesting that events on a time scale of ∼10 ms can be detected (**Fig. S2** in [18]).

### Kinetic analysis of DNA buckling

To quantify the extension time traces, we separate the extensions into two states by thresholding [26]. Upon stepwise increasing *ΔLK*, the time spent in the buckled state systematically increases at the expense of the population of the extended state (**Fig. 2a**). We analyze our experimental data using a two-state model [11,26], where the energy difference between the pre- and post-buckling states is related to *ΔLK*and the probability to be in the post-buckling state *P*_post_ is given by

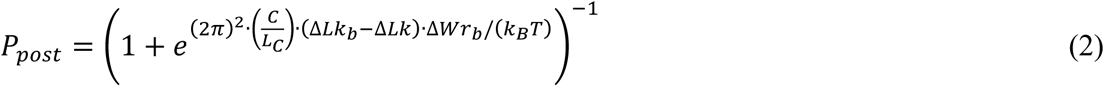

where *C* is the effective torsional stiffness (approximated in the Moroz-Nelson model [27] or using data from direct torque measurements [28] see [26]), *L*_*C*_ the contour length, *ΔLK*_*b*_ the buckling point, i.e. the number of applied turns for which *P*_pre=_ *P*_post_, Δ*Wr*_*b*_ *= Wr*_*post*_ *- Wr*_*pre*_ the number of turns converted from twist to writhe during buckling, *K*_*B*_ the Boltzmann constant and *T* the temperature. Fitting of Eqn. 2 to the experimentally observed *P*_*post*_ as a function of *ΔLK*yields *ΔLK*_*b*_ and *ΔWr*_*b*_ (**Fig. 2b**). Over the force range investigated, *ΔWr*_*b*_ remains essentially constant (**Fig. 2c**), in agreement with previous experimental results for both DNA [11] and RNA [15]. We find that *ΔWr*_*b*_ increases by ∼33% on increasing the monovalent salt concentration from 100 to 320 mM, again in quantitative agreement with previous findings [11,15].

**Figure 2.**
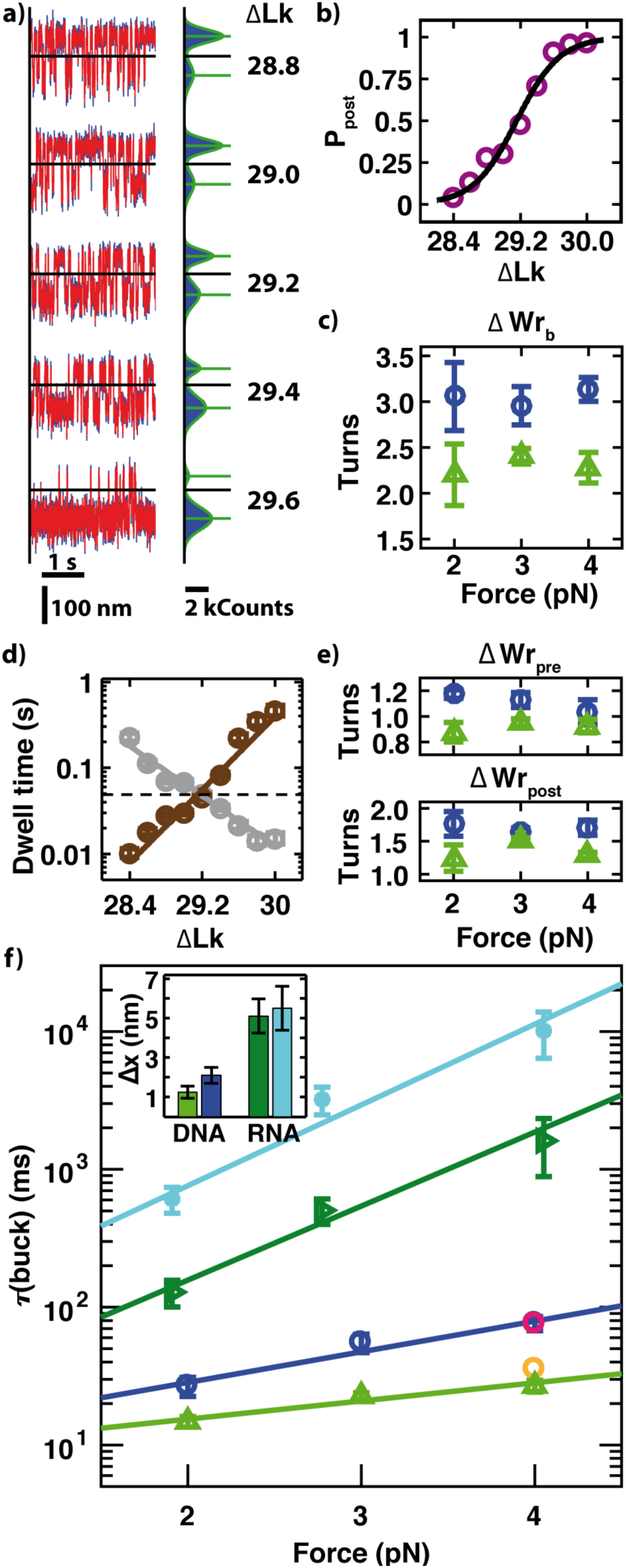
Quantification of DNA buckling dynamics. (a) Extension time traces as a function of *ΔLK*(Raw data 1000 Hz, blue; smoothed data 333 Hz, red). Black lines indicate thresholds for analysis. Histograms on the right are based on raw data and fitted by a double Gaussian. Horizontal green lines depict the mean extension of pre- and post-buckling state, respectively. (b) Fraction in the post-buckling state vs. *ΔLK*and fit of Eqn. 2 (solid line) at 3 pN and 320 mM NaCl. (c) Salt- and force dependence of *ΔWr*_*b*_ (green triangles 100 mM NaCl, blue circle 320 mM NaCl). (d) Mean dwell time for the pre- and post-buckling states vs. *ΔLK*and fits of Eqn. 3 (solid lines). The dashed line indicates the fitted buckling time *τ*_buck_. (e) Distances to the transition state |Δ*Wr*_pre_| and |Δ*Wr*_post_| (same color code as in c). (f) Buckling times vs. force for DNA (bottom two data sets; same color code as panel c) and RNA (top two data sets; dark green right-pointing triangle data are for 100 mM and cyan asterisk data for 320 mM NaCl) for different salt concentrations and exponential fits (Eqn. 4; solid lines). Magenta and orange circles for 4 pN, 320 mM NaCl taken from Ref [11] for 10.9 and 1.9 kbp DNA, respectively. Inset: Distances to transition state Δx for DNA and RNA. All RNA data taken from [15].

Having characterized the equilibrium properties of the buckling transition, we now focus on its dynamics, by analyzing the dwell times in the DNA extension traces. At each imposed *ΔLK*, the dwell time distributions for both the pre- and post-buckling state are well described by single exponential fits, which yield the mean lifetimes *τ*_pre_ and *τ*_post_ (**Fig. S3** in [18]). The lifetimes *τ*_pre_ and *τ*_post_ follow an Arrhenius relationship with an exponential dependence on *ΔLK*(**Fig. 2d**) [11]:

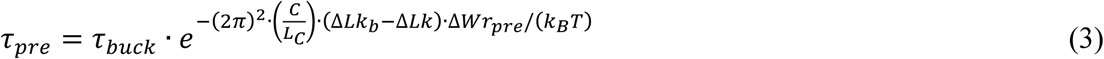

where *τ*_buck_ is the lifetime at the midpoint of the buckling transition *ΔLK*_*b*_, and *ΔWr*_*pre*_ is the change in writhe from the pre-buckling state to the transition state. We used a similar expression for *τ*_post_ with *ΔWr*_*pre*_ replaced by *-ΔWr*_*post*_, the rotational distance to the transition state from the post-buckling state (**Fig. 1c**).

Fits of Eqn. 3 to *τ*_pre_ and *τ*_post_ were used to determine the characteristic buckling time *τ*_buck_ as well as *ΔWr*_*pre*_ and *ΔWr*_*post*_ (**Fig. 2e**). The buckling time *τ*_buck_ increases with bead size, consistent with a model where the bead fluctuations are transmitted through the DNA molecule [29] (**Fig. S4** in [18]). For a fixed bead size, previous work [11] found a weak dependence of the characteristic buckling time on DNA length (comparing 1.9 and 10.9 kbp DNA gave a difference of 2-fold in the buckling times); our data for 7.9 kbp DNA under the same conditions falls between the previous measured buckling times, as would be expected for the intermediate length (**Fig. 2f,** differently colored points at 4 pN, 320 mM). For a fixed bead size and DNA length, the characteristic buckling time τ_buck_ is strongly dependent on force *F*, and is well described by an exponential model (solid lines in **Fig. 2f**, reduced *χ*^*2*^ *=* 0.4 and 0.3 for 100 and 320 mM NaCl, respectively):

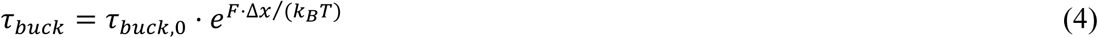

with *Δx* the distance to the transition state and τ_buck,0_ the buckling time in the absence of applied force. From a fit of Eqn. 4, we obtain τ_buck,0_ for DNA to be 8 ± 2 ms and 10 ± 3 ms (values and standard errors from the exponential fit, **Fig. 2f**) for 100 and 320 mM NaCl, respectively. These values agree with one another, within experimental error, and are close to the extrapolated buckling times at zero force for RNA [15] (τ_buck,0_ = 13 ± 7 ms and 52 ± 38 ms at 100 mM and 320 mM NaCl, respectively). The large differences between the buckling times τ_buck_ for DNA and RNA under tension, can primarily be attributed to differences in *Δx*, which is much smaller for DNA than for RNA under otherwise identical conditions (**Fig. 2f inset**), supporting our previous hypothesis that the striking difference in the buckling dynamic between DNA and RNA is mostly driven by differences in the transition state. We note that supercoil nucleation does not merely depend on force, but on torque as well (the different points in Fig. 2f are not only at different forces, but also at different torques, since the buckling point *ΔLk*_*b*_ shifts significantly with applied force). Therefore, *Δx* is a value that quantifies the position of the energy barrier in a simplified 1D representation of the energy landscape. Its value does not directly reflect a physical position of the transition state, but should rather be interpreted as a characteristic length that describes the cooperativity of the transition. A full description of the transition requires considering the energy landscape along rotational (twisting, writhing, and linking) and extension (DNA end-to-end distance) degrees of freedom, see below.

To quantify the energy landscape of supercoil nucleation along the rotational degree of freedom, we first determined the distances to the transition state from the pre-buckling and post-buckling states *ΔW*_*pre*_ and *ΔWr*_*post*_ from fits of Eqn. 3 (**Fig. 2d**). Both *ΔW*_*pre*_ and *ΔWr*_*post*_ change systematically with salt concentration, but remain approximately constant with increasing force (**Fig. 2e**) and bead size (**Fig. S4** in [18]). Notably, *ΔWr*_*post*_ and *ΔWr*_*pre*_ add up to the measured value for *ΔWr*_*!*_, within experimental error (**Fig. S5** in [18]), as would be expected, since they are measured along the same coordinate. The ratio *ΔWr*_*pre*_*/ΔWr*_*post*_ is independent of force and ionic strength, within experimental error, and suggests the transition state along the writhing degree of freedom to be closer to the pre-buckling state than the post-buckling plectonemic state (*|ΔWr*_*pre*_*/ΔWr*_*post*_| = 0.68 ± 0.05 and 0.65 ± 0.03 for 100 mM and 320 mM NaCl, respectively). Since the transition occurs at a constant *ΔLk*, the measured ratio of *|ΔWr*_*pre*_*/ΔWr*_*post*_| implies that the transition state in the twisting degree of freedom is closer to the post-buckling state than the pre-buckling state.

### Energy landscape reconstruction at buckling equilibrium

To obtain a full quantitative description of the buckling transition and to account for its mutual dependence on extension and rotation, we reconstructed the 2D free energy landscape *ΔG(z, ΔLK)*. At a given *ΔLK, ΔG z = -K*_*B*_*T. ln[p(z)]* with the probability density *p(z)* [30]. To account for the effect of the force probe, we deconvolved the extension histogram of the DNA tether with the setup response function *S(z)* [30-33]. For the deconvolution procedure, we used the bead fluctuations of torsional unconstrained DNA molecules at the same force and buffer conditions as the instrument response function (see [18]). The deconvolution sharpens the extension histogram (**Fig. 3a**), which is then converted to the corresponding 1D energy landscape to enable the analysis of the buckling transition along the extension coordinate. We compared the energy landscape obtained by deconvolution to the energy barrier reconstructed by a different approach, based on the splitting probability [34,35] with no need for deconvolution [33], and obtained very good agreement between the two methods (see **Supporting Materials** [18] and **Supplementary Fig. S6**). Finally, by assembling a series of 1D extension energy landscapes while systematically varying *ΔLK*, we reconstruct the full 2D extension-linking number energy landscape **(Fig. 4b)**.

**Figure 3.**
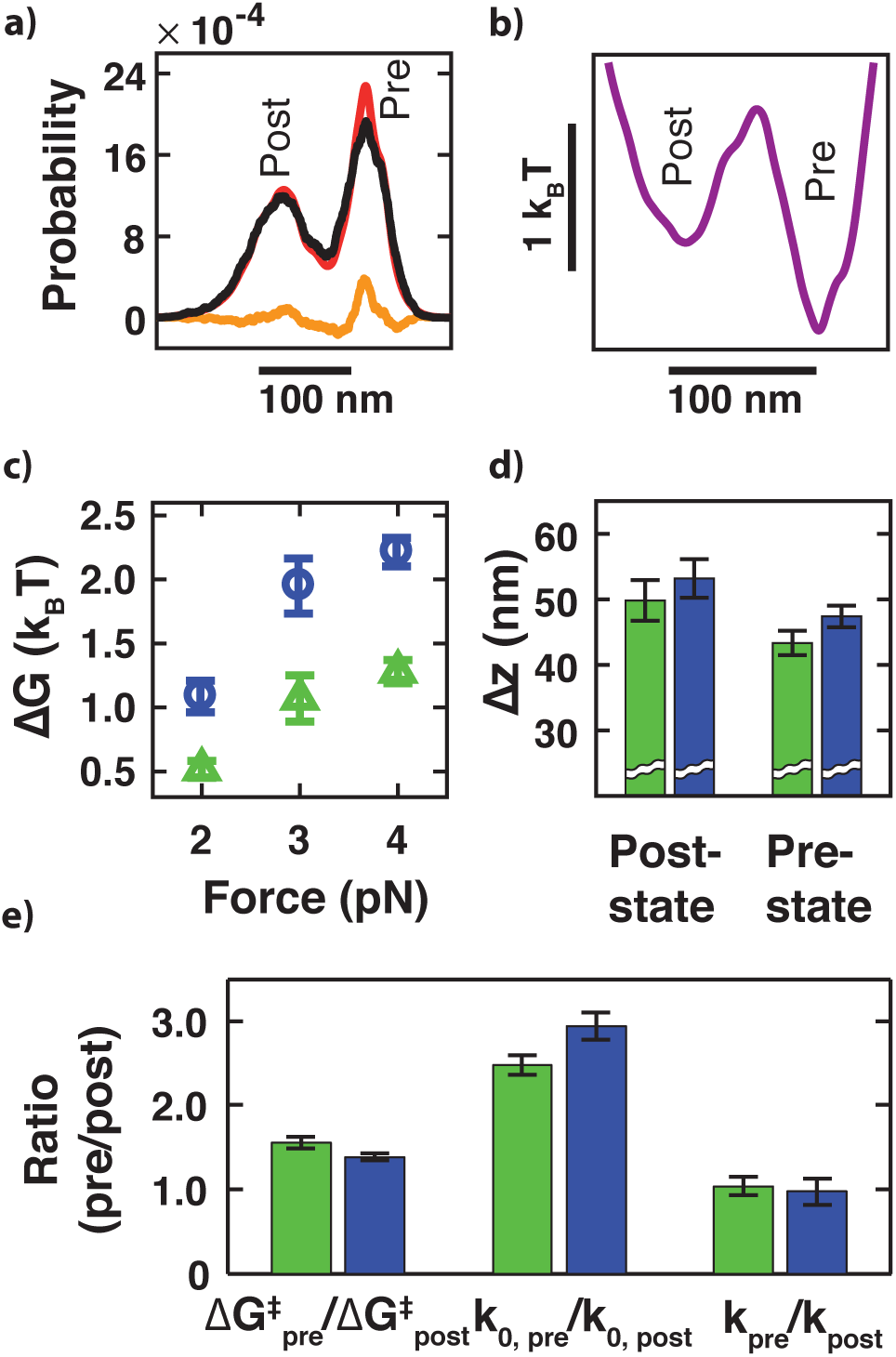
Reconstruction and quantification of the energy landscape. (a) Deconvolution of extension histograms. Extension histogram before (black) and after (red) deconvolution with the setup response function and difference due to deconvolution (orange). (b) Reconstructed energy landscape computed as ΔG(z) = –k_B_T·ln[p(z)] from the deconvolved probability density p(z). (c) Mean value of the energy barrier at *ΔLK*_*b*_ of pre- and post buckling state vs. force (color code for panels c-e: 100 mM NaCl green and 320 mM NaCl blue). (d) Distances to the energy barrier along the extension coordinate (absolute values). (e) Comparison of post and pre-buckling state parameters at *ΔLK*_*b*_: The energy barrier for the pre-buckling state 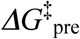 is larger than for the post buckling state 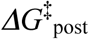 resulting in 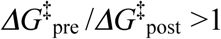. The ratio of the attempt frequency of the bead to cross the energy barrier of the pre-buckling state by the post-buckling state k_0,pre_/k_0,post_ determined by the ratio of the curvature of the energy landscape is larger than 1. The overall rate to cross the energy barrier k_pre_/k_post_ ≈ 1 in line with equilibrium at *ΔLK*_*b*_.

**Figure 4.**
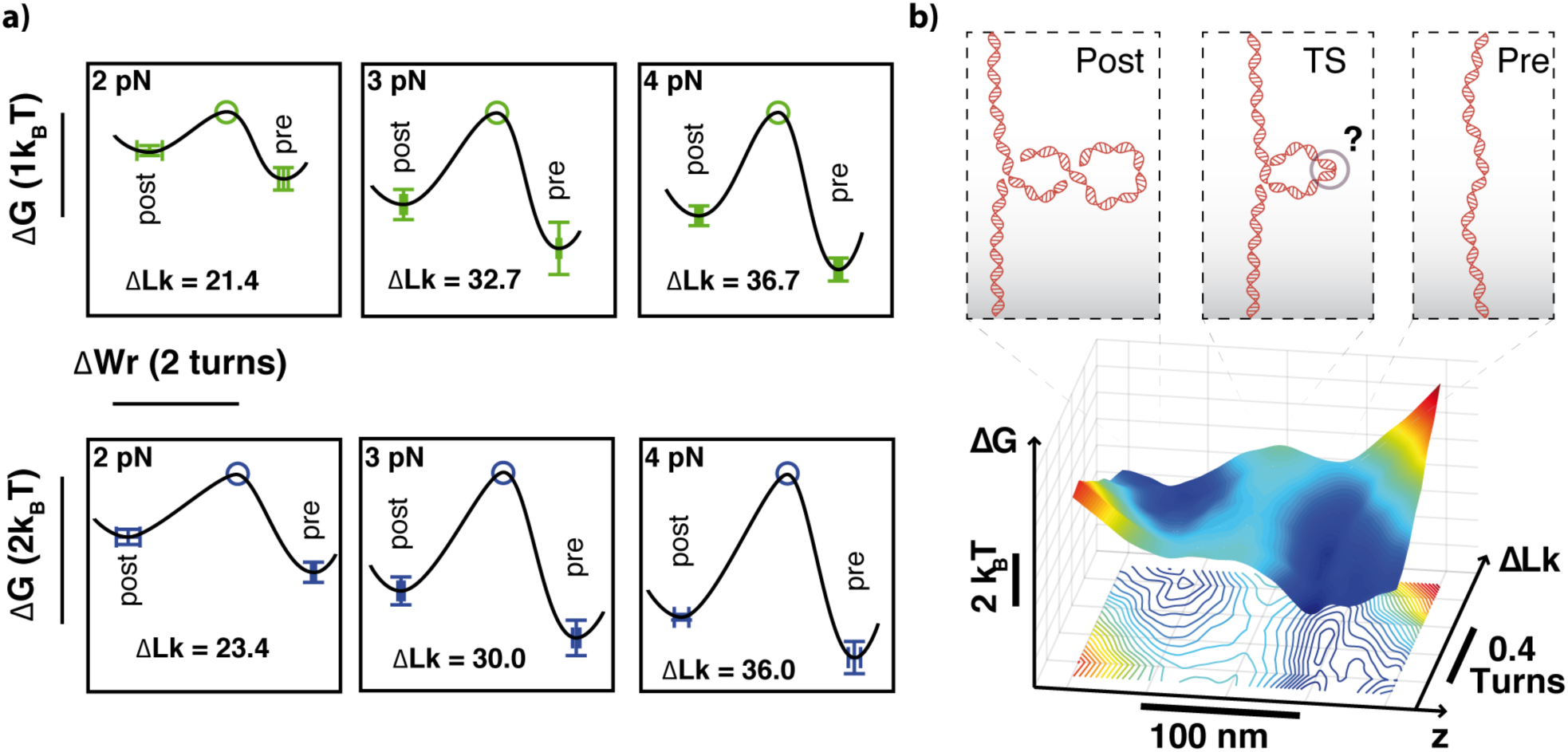
Energy landscape of the DNA buckling transition. (a) Quantitative energy landscape along *ΔWr* for different forces and salt concentrations (upper row 100 mM NaCl, lower row 320 mM NaCl). Green and blue markers are determined from the analysis described in the main text. The energy landscape flattens for lower forces. Since the analysis only provides energy differences, the top of the barriers are set to the equal height for ease of comparison. (b) A full 2D extension and rotation energy landscape was obtained by subsequently constructing 1D extension energy landscapes (as in Fig. 3) while systematically changing *ΔLK*(in steps of 0.1 turns). The iso-energy lines in the projection of the landscape into 2D have a spacing of 0.15 k_B_T. The asymmetry of the pre- and post-buckling energy wells is apparent.

We analyze the reconstructed energy landscape focusing on the extension coordinate at *ΔLK*_*b*_, i.e. at the point where the forward and backward rates are equal [26]. At *Δ LK*_*b*_, the distances to the transition state along the extension coordinate from the pre- and post-buckling state minima, *Δz*_*pre*_ and *Δz*_*post*_, were found to be dissimilar, independent of force (**Fig. S7** in [18]). The values suggest that the transition state along the extension coordinate is closer to the pre- than to the post-buckling state **(Fig. 3d)**. Notably, *Δz*_*pre*_ and *Δz*_*post*_ add up to the total jump size as determined directly from extension distributions (**Fig. S8** in [18]). At *ΔLK*_*b*_, the reconstructed energy landscape (**Fig. 3b** and **Fig. S8b** in [18]) shows a broader energy minimum for the post-buckling state compared to the pre-buckling state, corresponding to a larger conformational space after buckling. The broader energy minimum after buckling corresponds to a smaller curvature (**Fig. S9** and **S10** in [18]) and, applying Kramers theory [31,36,37], to a smaller attempt rate *k*_*0*_ for barrier crossing at buckling equilibrium compared to the pre-buckling state. Since the forward and backward rates are identical at *ΔLK*_*b*_, the difference in *k*_*0*_ is compensated by different barrier heights 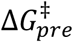 and 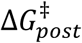 (**Fig. S9c,d** and **S7c,f** in[18]) measured from either side of the transition state. We conclude that the energy barrier, measured along the extension coordinate, is significantly asymmetric and steeper for buckling as for plectoneme removal.

The reconstructed energy landscape enable, in addition, the calculation of diffusion coefficients for barrier crossing, again from Kramer’s theory (see e.g. Equation 10 from Ref. [31]). We find diffusion coefficients *D* ≈ 10^−15^ m^2^/s, whereby the diffusion constants for the pre- and post-buckling wells are essentially within error, but slightly force and salt dependent. Values of *D* ≈ 10^−15^ m^2^/s are in the same order of magnitude as reported for protein relaxation times and refolding landscapes [38,39] and smaller than DNA hairpin diffusion coefficients [31]. The estimated diffusion coefficient of 10^−15^ m^2^/s implies significantly slower diffusion than what would be estimated for purely translational diffusion of DNA with length corresponding to the loop size, which is in the range of ∼10^−11^-10^−12^ m^2^/s [40]. Thus, diffusive barrier crossing is several orders of magnitude slower as compared to simple translational diffusion, which in turn implies substantial internal friction in the DNA as it evolves to the transition state.

The energy landscape reconstruction is not limited to the extension coordinate (**Fig. 3**); we can also quantitative describe the energy landscapes along the rotational degree of freedom (**Fig. 4a**). Using the values for *ΔWr*_*post*_ and *ΔWr*_*pre*_ (**Fig. 2e**) to determine the relative position along the writhe coordinate and the free energy barrier heights from the energy landscape (**Supplementary Figure S7c,f**), we obtain a free energy landscape along the *ΔWr* coordinate at fixed *ΔLK*_*b*_ that again reveals an asymmetrical distance to the transition state. The energy barrier flattens with a decrease in force (see also **Supplementary Fig. S7**) in line with previous observations for force dependent energy barriers [41].

## CONCLUSION

In summary, taking advantage of the ability of MT to control both the applied force and twist, we have reconstructed the full 2D energy landscape along the extension and twist degrees of freedom. Along the rotational degree of freedom the energy landscape is asymmetric, with *ΔWr*_*pre*_ close to unity, and roughly 30% smaller as compared to *ΔWr*_*post*_. Likewise, the energy landscape along the extension coordinate exhibits significant asymmetry, with the distance from the extended to the transition state *Δz*_*pre*_, approximately 15 % smaller than the distance from the buckled to the transition state *Δz*_*post*_. Combining both results, we conclude that the transition state is a small, highly twisted, single loop.

We hypothesize that the strong bending and twisting deformations in the transition state for DNA buckling lead to a breakdown of isotropic elasticity and result in the formation of a kinked loop [42]. A breakdown of harmonic elasticity for DNA could explain the finding from the force dependence of τ_buck_ that the buckling transition exhibits a three- to four- fold steeper energy barrier for DNA as compared to RNA (**Fig. 2f**). Barrier steepness quantifies the cooperativity of molecular rearrangements to achieve the transition state geometry; since the (harmonic) elasticity properties of both nucleic acids are roughly the same, it is plausible that the transition involves the breakdown of DNA harmonic elasticity. This hypothesis is further supported by the observation that factors that destabilize double-stranded DNA, notably glycerol or low salt concentrations **(Fig. S11** and **S12** in **[18])**, increase the rate of buckling, consistent with a lower transition state energy barrier. Disruption of DNA base pairing upon negative supercoiling under stretching forces has been clearly established previously [43,44]. Our hypothesis of kinking and local disruption of base pairing is in line with biochemical and structural experimental results [45] and with all-atom molecular dynamics simulations of small DNA circles [46] that indicate the formation of local kinks also upon positive supercoiling.

Our findings suggest that local defects [11,47], e.g. introduced by DNA damage or protein-binding [48], would enhance the rate of supercoil nucleation by transition state stabilization, and help positioning plectonemes. The rates of supercoil location hopping, previously determined using identical beads [13], are similar to the rates of supercoil nucleation determined in this study, which strongly suggests that long-range communication along DNA is rate-limited by supercoil nucleation. In summary, the quantitative framework presented here will enable making testable predictions of DNA topology-mediated regulatory dynamics and provides a critical baseline for models of DNA dynamics *in vivo*. In addition, our work highlights the necessity to decouple the energy landscape of supercoil nucleation along both extension and rotational degrees of freedom, and demonstrates how high-speed magnetic tweezers experiments allow reconstruction of full 2D free energy landscapes, which opens up exciting possibilities to extend the more commonly used 1D free energy description of macromolecular transitions [49] into multiple dimensions [50].

## Acknowledgements

We thank Samuel Stubhan for help with initial measurements, Daniel Burnham for sharing code for the bead-DNA simulations, Enrico Carlon and Stefanos Nomidis for stimulating discussions, Angelika Kardinal and Thomas Nicolaus for help with sample preparation, and the German Research Foundation (DFG) through SFB 863, KU Leuven through IDO, and FWO Flanders, for funding.

## REFERENCES

[1] L. F. Liu and J. C. Wang, Proc Natl Acad Sci U S A 84, 7024 (1987).

[2] N. R. Cozzarelli, G. J. Cost, M. Nollmann, T. Viard, and J. E. Stray, Nature reviews. Molecular cell biology 7, 580 (2006).

[3] D. A. Koster, A. Crut, S. Shuman, M. A. Bjornsti, and N. H. Dekker, Cell 142, 519 (2010).

[4] T. J. Stevens et al., Nature 544, 59 (2017).

[5] F. B. Fuller, Proc Natl Acad Sci U S A 75, 3557 (1978).

[6] J. H. White, American Journal of Mathematics 91, 693 (1969).

[7] A. V. Vologodskii and N. R. Cozzarelli, Annual review of biophysics and biomolecular structure 23, 609 (1994).

[8] T. Schlick, Current opinion in structural biology 5, 245 (1995).

[9] J. Vinograd, J. Lebowitz, R. Radloff, R. Watson, and P. Laipis, Proc Natl Acad Sci U S A 53, 1104 (1965).

[10] T. R. Strick, J. F. Allemand, D. Bensimon, A. Bensimon, and V. Croquette, Science 271, 1835 (1996).

[11] H. Brutzer, N. Luzzietti, D. Klaue, and R. Seidel, Biophys J 98, 1267 (2010).

[12] J. Dekker and L. Mirny, Cell 164, 1110 (2016).

[13] M. T. J. van Loenhout, M. V. de Grunt, and C. Dekker, Science 338, 94 (2012).

[14] A. Crut, D. A. Koster, R. Seidel, C. H. Wiggins, and N. H. Dekker, Proc Natl Acad Sci U S A 104, 11957 (2007).

[15] J. Lipfert et al., Proc Natl Acad Sci U S A 111, 15408 (2014).

[16] S. Forth, C. Deufel, M. Y. Sheinin, B. Daniels, J. P. Sethna, and M. D. Wang, Physical review letters 100, 148301 (2008).

[17] B. C. Daniels and J. P. Sethna, Physical review. E, Statistical, nonlinear, and soft matter physics 83, 041924 (2011).

[18] See Supplemental Material at [URL will be inserted by publisher] for material and methods as well as detailed analysis.

[19] L. Postow, C. D. Hardy, J. Arsuaga, and N. R. Cozzarelli, Genes & development 18, 1766 (2004).

[20] J. Lipfert, X. Hao, and N. H. Dekker, Biophys J 96, 5040 (2009).

[21] B. M. Lansdorp and O. A. Saleh, The Review of scientific instruments 83, 025115 (2012).

[22] J. F. Allemand, D. Bensimon, R. Lavery, and V. Croquette, Proceedings of the National Academy of Sciences 95, 14152 (1998).

[23] B. M. Lansdorp, S. J. Tabrizi, A. Dittmore, and O. A. Saleh, The Review of scientific instruments 84, 044301 (2013).

[24] A. Huhle, D. Klaue, H. Brutzer, P. Daldrop, S. Joo, O. Otto, U. F. Keyser, and R. Seidel, Nature communications 6, 5885 (2015).

[25] D. Dulin, T. J. Cui, J. Cnossen, M. W. Docter, J. Lipfert, and N. H. Dekker, Biophys J 109, 2113 (2015).

[26] P. Daldrop, H. Brutzer, A. Huhle, D. J. Kauert, and R. Seidel, Biophysical Journal 108, 2550 (2015).

[27] J. D. Moroz and P. Nelson, Proc Natl Acad Sci U S A 94, 14418 (1997).

[28] J. Lipfert, J. W. Kerssemakers, T. Jager, and N. H. Dekker, Nature methods 7, 977 (2010).

[29] H. Bai, J. E. Kath, F. M. Zorgiebel, M. Sun, P. Ghosh, G. F. Hatfull, N. D. Grindley, and J. F. Marko, Proc Natl Acad Sci U S A 109, 16546 (2012).

[30] M. T. Woodside, P. C. Anthony, W. M. Behnke-Parks, K. Larizadeh, D. Herschlag, and S. M. Block, Science 314, 1001 (2006).

[31] M. T. Woodside and S. M. Block, Annual review of biophysics 43, 19 (2014).

[32] P. Jansson, A., Deconvolution of images and spectra (2nd ed.) (Academic Press, Inc., 1996).

[33] A. P. Manuel, J. Lambert, and M. T. Woodside, Proc Natl Acad Sci U S A 112, 7183 (2015).

[34] R. Du, V. S. Pande, A. Y. Grosberg, T. Tanaka, and E. S. Shakhnovich, The Journal of Chemical Physics 108, 334 (1998).

[35] J. D. Chodera and V. S. Pande, Physical review letters 107, 098102 (2011).

[36] H. A. Kramers, Physica 7, 284 (1940).

[37] P. Hänggi, P. Talkner, and M. Borkovec, Reviews of Modern Physics 62, 251 (1990).

[38] R. Berkovich, R. I. Hermans, I. Popa, G. Stirnemann, S. Garcia-Manyes, B. J. Berne, and J. M. Fernandez, Proceedings of the National Academy of Sciences 109, 14416 (2012).

[39] H. Lannon, J. S. Haghpanah, J. K. Montclare, E. Vanden-Eijnden, and J. Brujic, Physical review letters 110, 128301 (2013).

[40] E. Stellwagen and N. C. Stellwagen, Analytical chemistry 87, 9042 (2015).

[41] B. Sudhanshu, S. Mihardja, E. F. Koslover, S. Mehraeen, C. Bustamante, and A. J. Spakowitz, Proc Natl Acad Sci U S A 108, 1885 (2011).

[42] T. A. Lionberger, D. Demurtas, G. Witz, J. Dorier, T. Lillian, E. Meyhofer, and A. Stasiak, Nucleic acids research 39, 9820 (2011).

[43] T. R. Strick, V. Croquette, and D. Bensimon, Proc Natl Acad Sci U S A 95, 10579 (1998).

[44] C. Matek, T. E. Ouldridge, J. P. Doye, and A. A. Louis, Scientific reports 5, 7655 (2015).

[45] R. N. Irobalieva et al., Nature communications 6, 8440 (2015).

[46] J. S. Mitchell, C. A. Laughton, and S. A. Harris, Nucleic acids research 39, 3928 (2011).

[47] A. Dittmore, S. Brahmachari, Y. Takagi, J. F. Marko, and K. C. Neuman, Physical review letters 119, 147801 (2017).

[48] J. F. Marko and S. Neukirch, Physical review. E, Statistical, nonlinear, and soft matter physics 85, 011908 (2012).

[49] O. K. Dudko, G. Hummer, and A. Szabo, Proceedings of the National Academy of Sciences 105, 15755 (2008).

[50] Y. Suzuki and O. K. Dudko, Physical review letters 104, 048101 (2010).

